# Detecting stabilizing, directional, and disruptive patterns of anthropogenic species loss with general models of nonrandom extinction

**DOI:** 10.1101/2022.09.11.507476

**Authors:** Nicholas A. Huron, S. Blair Hedges, Matthew R. Helmus

## Abstract

The selective landscape that gave rise to Earth’s species has shifted in the Anthropocene. Humans have accelerated extinction pressures, making efforts to detect general non-random patterns of extinction increasingly important. Much research has focused on detecting which traits make some species more likely to go extinct, such as large body size and slow reproductive rate in animals, limited dispersal in vascular plants, and narrow habitat requirements in cacti. However, general models for such non-random extinction are lacking. Here, we adapt the three general models of natural selection to classify non-random extinction as directional, disruptive, or stabilizing extinction. We develop a quantitative method for testing which general extinction model best describes observed data and apply it to the Caribbean lizard genus *Leiocephalus* as a case study. We surveyed the literature for recorded last occurrence for extinct and threat status for extant species. Eight species have gone extinct and ten are predicted to go extinct soon. Past extinctions in *Leiocephalus* showed directional extinction of large bodied species, while future-predicted extinctions exhibited a more complex extinction model similar to both random and stabilizing extinction with respect to body size. Similarly, future-predicted extinctions exhibited stabilizing extinction with respect to limb and tail lengths. Lizards with either very long or very short appendages are most likely to go extinct in the future. This shift from directional to stabilizing extinction for *Leiocephalus* is consistent with hunting, introduced predators, and habitat loss that first increased extinction pressure on the largest species and then extinction pressure on species that deviate from an adaptive peak centered on a generalist ground-lizard body plan. As adaptive optima shift in the Anthropocene, general models of non-random extinction are essential to developing a mature strategy for future successful conservation efforts.

## Introduction

Extinction can be evaluated with respect to traits of the extinct species in comparison to extant species (Hooper et al. 2005; Díaz et al. 2007; Cardinale et al. 2012; Tilman et al. 2014). When species go extinct within a clade, the loss may be broadly considered random or non-random for a particular trait. Random extinction is a stochastic process to describe background extinction rates that occurs irrespective of species traits, while non-random extinction is loss that deviates from random with respect to measured traits (Purvis et al. 2000; Raffaelli 2004). Non-random extinction is often used to connect extinction to specific and often anthropogenic impacts (Brook and Alroy 2017). For animals, common attributes that are non-random indicators of extinction include life history traits like large body size and slow reproductive rate, small population size or geographic range, high sensitivity to environmental stress that is exacerbated by humans, and phylogenetic distinctiveness (Purvis et al. 2000; Cardillo 2003; Raffaelli 2004; Vamosi and Wilson 2008; Yessoufou and Davies 2016). Non-random extinction of species with certain traits can be especially concerning when those traits can be linked to ecosystem functions (Cardinale et al. 2012; Mouillot et al. 2013), a phenomenon that has been reported for several systems (Brook et al. 2008; Dirzo et al. 2014; Chichorro et al. 2019).

Efforts to identify patterns of non-random extinction, also termed extinction selectivity (Raup 1994), have first focused on broad taxonomic categories and easily measured traits to identify extinction-prone taxa (Grenyer et al. 2006; Brook et al. 2008; Barnosky et al. 2011; Forest et al. 2015; Lyons et al. 2016; Yessoufou and Davies 2016; Chichorro et al. 2020)., the advent of large molecular phylogenetic datasets has made quantifying extinction and its risk with phylogenetic relationships more tractable than ever before (Isaac et al. 2007; Davies and Yessoufou 2013; Forest et al. 2015; Yessoufou and Davies 2016; Maliet et al. 2018). Such advances have prompted development of modeling methods that consider phylogeny and traits jointly to identify non-random extinction (e.g., BiSSE, QuaSSE, and related SSE models; Maddison et al. 2007; FitzJohn 2010), but these methods have their limitations. For example, phylogenies are useful for measuring diversification (speciation and extinction), but it may be impossible to accurately resolve reliable extinction records for empirical data with them, since they must identify when species are no longer present in a phylogeny (Purvis 2008; FitzJohn 2010). Moreover, the number of species that can be evaluated using modern molecular phylogenetic methods is increasing, but there are many species for which data are still unavailable (Mora et al. 2011; Yessoufou and Davies 2016; Lewin et al. 2022). Therefore, it is wise to seek other methods alongside phylogenetic-based ones for identifying when extinctions are non-random for particular characteristics (Purvis 2008), especially when they correspond with contributions to ecosystem function (Forest et al. 2015; Hull 2015). Although some studies to date test for non-random vs. random anthropogenic extinction, few focus on classifying observed non-random extinction beyond identifying if an extreme trait state seems related to loss. This is especially clear in reviews of empirical extinction selectivity (e.g., McKinney 1997; McKinney and Lockwood 1999; Brook et al. 2008; Barnosky et al. 2011; Hull 2015; Yessoufou and Davies 2016; Brook and Alroy 2017), which are often reduced to lists of traits associated with extinction. Oversimplification of extinction selectivity in turn seems to limit how it is modeled (e.g., Gross and Cardinale 2005 but see Ciampaglio et al. 2001; Korn et al. 2013; Puttick et al. 2020 for slightly improved approaches). Although methods for describing non-random extinction increasingly consider different extinction selectivity patterns, they lack a cohesive set of models to evaluate extinctions against. In contrast, natural selection with respect to traits has long been classified into three separate models. To address this gap we develop a method for evaluating how extinction for a clade affects diversity of a trait over time.

To classify non-random extinction, we propose to follow a similar classification scheme to the general natural selection models from population genetics (Rueffler et al. 2006). Specifically, if one considers an adaptive landscape with one or more peaks corresponding to maximum species viability, then three general selection models emerge: directional selection can lead species to shift trait frequencies towards a peak from its base, disruptive selection can lead species out of adaptive minima in rugged landscapes, and stabilizing selection keeps species near a peak once they approach it (Arnold et al. 2001; Pfaender et al. 2016). Similarly, when one considers non-random deviations from random extinction with respect to species traits, three similarly-named possible models arise: (1) directional, (2) disruptive, or (3) stabilizing extinction (Figure 1). Directional extinction, whereby loss is greater for species at one trait distribution extreme, is the most commonly reported form of non-random extinction in literature (e.g., large-bodied animals; Pregill 1986; Cardillo 2003; Cardillo et al. 2005; Kemp and Hadly 2015; Yessoufou and Davies 2016). The second model, disruptive extinction, occurs when species with intermediate trait values (those closest to the mean) experience greater rates of extinction than the extremes. Although the location of extinction pressure differs from directional extinction, the idea that a single portion of the distribution is preferentially lost is common between the two. Disruptive extinction is elusive in the literature but has been reported for body mass in Australian (Woinarski et al. 2017) and New Zealand (Duncan and Blackburn 2004) bird species and dispersal ability in butterflies (Thomas 2000). Lastly, stabilizing extinction describes the case in which two loci of extinction pressure exist, whereby species with extreme trait values are preferentially lost (e.g., predicted threat for body size in multiple vertebrate taxa; Ripple et al. 2017; Leclerc et al. 2018).

**Figure 1:**
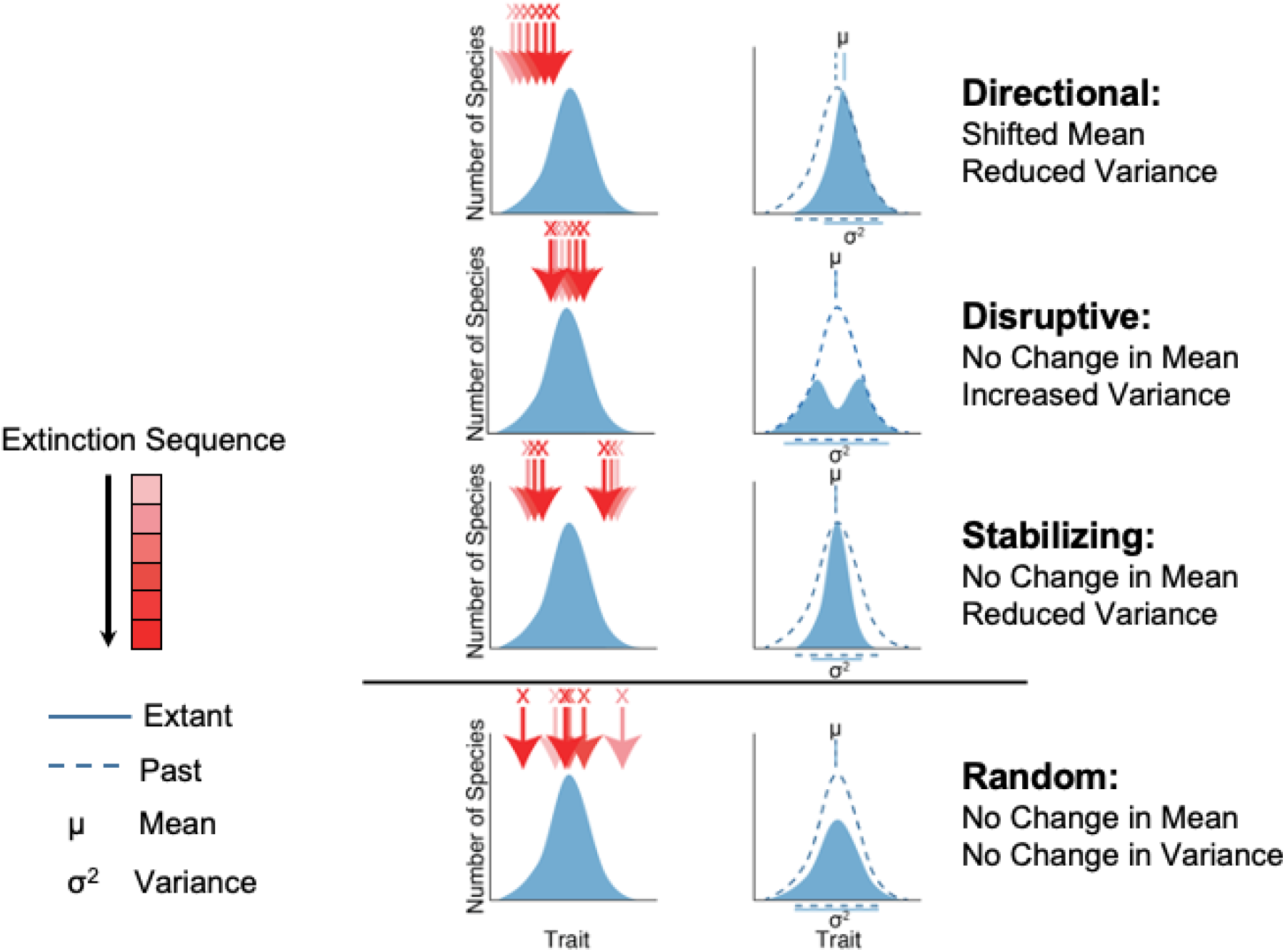
Extinction models conceptualized. Extinction for a particular trait for a clade of interest can be random or non-random based on changes to the clade-wide trait mean and variance as species are lost. In the first column, extinction annotations (red arrows and x’s) show the localized area(s) of the distribution where loss occurs for a particular extinction model (proceeding in increasing color intensity). The second column shows the expected change in the absolute frequency distribution that follows each extinction model and includes estimates of μ and σ^2^ for the past (dashed) and extant (solid) trait distributions.

### Three general models of nonrandom extinction

Based on extinction model expectations, one can predict how a trait distribution for a clade will change as species are lost. First, suppose a clade of *n* species (*i* = 1, … *n*), each made up of many individuals that all have a continuous trait, *x*. For this scenario, we ignore intraspecific trait variation and only focus on the intraspecific mean for *x* for each species *i*, 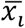, and how it relates to variation in extinction probability among species within the clade. Values of 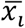 follow a Gaussian distribution with a clade mean, μ, standardized to zero, and clade variance, σ^2^, standardized to one for simplicity. As species go extinct, μ and σ^2^ can change, depending on 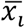 of lost species. If loss is random, the expected values of μ and σ^2^ will not change. Random extinction follows a uniform probability distribution that is independent of species trait values, which leads to the same expected clade mean and variance (Δμ = 0, Δσ^2^ = 0) following a set of sequential extinctions that have occured over time (i.e., an extinction sequence, *i* = 1, …, *e* total extinctions). In contrast, non-random extinction produces different Δμ and Δσ^2^ depending on the model of extinction it follows (Figure 1). Directional extinction exhibits changes in both μ and σ^2^ due to disproportionate loss of species from one of the tails of the distribution. The sign of Δμ is negative if loss occurs from the right tail (species with the largest trait values are lost) or positive if loss occurs from the left tail (species with the smallest trait values are lost). Under both directional extinction sequences, Δσ^2^ is negative. Disruptive and stabilizing extinction exhibit no expected change to the clade mean (Δμ = 0), but opposite expected Δσ^2^ signs.

Disruptive extinction is characterized by loss of species near the center of the distribution (those closest to μ), leading to a positive Δσ^2^, whereas stabilizing extinction corresponds with loss of species with extreme values (those farthest from μ), resulting in a negative Δσ^2^. Therefore, methods that observe Δμ and Δσ^2^ for an extinction sequence should be able to distinguish the three general models of non-random extinction from one another and from non-random extinction.

To test the three general non-random extinction models and their corresponding predicted changes to a trait distribution across species in a clade, we employed an empirical case study. Specifically, we evaluated past and predicted-future extinction in a clade from the Caribbean Islands biodiversity hotspot (Myers et al. 2000; Mittermeier et al. 2004; Zachos and Habel 2011). Although Caribbean islands are biodiverse, reptiles in particular are one of the most speciose, yet imperiled vertebrate groups found there (Myers et al. 2000; Mittermeier et al. 2004; Ricklefs and Bermingham 2008; Böhm et al. 2013). Several taxa typify distinct Caribbean reptile diversity (e.g., the well-studied *Anolis* adaptive radiation; Losos 2009), but one particular genus is an exemplar for diversity, endemicity, and threat of extinction: *Leiocephalus*, or curly-tailed lizards (Mittermeier et al. 2004; Hedges 2022). Over half of described *Leiocephalus* are either extinct or threatened with extinction (25% and 34.4% of species, respectively; IUCN 2020). Proposed sources of past extinction include introduced mammal predators and selective foraging by humans while current threats also include habitat loss (Steadman et al. 1984; Powell et al. 1999; Daltry 2007; Borroto-Páez and Woods 2012; IUCN 2020). Kemp and Hadly (2015) reported in a study of Caribbean lizards that larger *Leiocephalus* species have gone extinct, thereby suggesting past directional extinction. We first articulated a method to test the three general models of non-random extinction and performed simulations to produce expected distributions under each model to assess observed patterns. We then demonstrated its practical implementation to investigate past and expected future extinctions for *Leiocephalus* morphological traits. We found that past and expected future extinctions provide evidence for different patterns of non-random extinction and thereafter discuss the implications of shifting non-random extinction.

## Methods

### Case Study

We evaluated which non-random extinction model best explained observed extinctions for the lizard genus, *Leiocephalus*, which comprises 32 described species from the Caribbean (Henderson and Powell 2009; Köhler et al. 2016; Hedges 2022). According to current IUCN threat assessments (IUCN 2020), 8 species are extinct, and of the 24 extant species, 13 species are least concern (LC), 1 is near threatened (NT), 2 are vulnerable (VU), 1 is endangered (EN), 5 are critically endangered (CR), and 2 are data deficient (DD). Since *Leiocephalus* has experienced changes in extinction pressures, we use this clade as an empirical case study to address if past extinctions and expected future extinctions follow the same extinction model.

#### Observed Extinction Sequence

For *Leiocephalus*, we used the most recently published species IUCN threat status (IUCN 2020). We assigned the two DD species based on the best available data: *Leiocephalus sixtoii* to VU (personal observation, SBH) and *Leiocephalus varius* to LC based on the lack of a prior record in the IUCN red list. For all analyses, we used these updated statuses to place species into three groups: extinct (EX species), threatened (all extant but not LC species), and least concern species (extant LC species). Dates were centered at 1950CE, which was set to 0 to ensure continuity between geologic time and dates that use CE across literature sources.

To estimate the past *Leiocephalus* extinction sequence, date ranges for extinct species were based on two possible approaches. If a species had an estimated epoch or age of last occurrence in the literature (Etheridge 1965, 1966; Pregill 1981, 1984), published start and end dates for corresponding epochs or ages were used. For all other extinct species, the year last seen, as recorded in current IUCN data (IUCN 2020), was used. IUCN data include both individual years and date ranges; the latter were used whenever possible. For all single year data, we treated that year as an extinction date midpoint for a range buffered on each side by 10 years.

For predicted-future extinction sequence, we used current IUCN threat status and red list categories criterion E (IUCN 2012) to extrapolate the future date (in years) of likely extinction. IUCN criterion E sets a probability of extinction within a particular number of years for CR (*P*_*extinct*_ ≥ 0.50 within ten years), EN (*P*_*extinct*_ ≥ 0.2 0 within twenty years), and VU (*P*_*extinct*_ ≥ 0.10 within one hundred years) species. Corresponding probabilities are not published for NT or LC species. Therefore, we followed Oliveira et al. (2020) for both categories (NT: *P*_*extinct*_ ≥ 0.01 within one hundred years; LC: *P*_*extinct*_ ≥ 0.0001 within one hundred years).

We solved for the date of extinction for *P*_*extinct* =_ 0.9*5* as extrapolated from criterion E probabilities by assuming that the probability of extinction and remaining extant over the same time interval sum to unity (1).

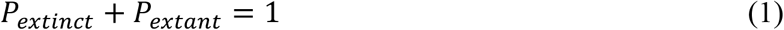

From this, we treated the probability of remaining extant as an exponential decay function (2):

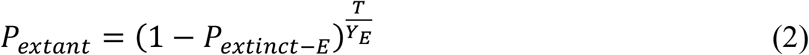

where *P*_*extinct*−*E*_ is the probability of extinction, *Y*_*E*_ describes the time interval in years for each category from criterion E in years, and *T* describes the time in years it takes to fulfill *P*_*extant*_ at the set value. By algebraic manipulation of (2), one can arrive at (3):

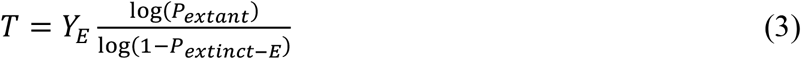

We set *P*_*extant*_ = 0.0*5* and calculated the value of *T* for all extant *Leiocephalus* based on red list categories. The resulting *T* values were treated as extinction date midpoints with a single year last seen (*T* ±10 years).

The exact year of extinctions is unknown for all the species, thus there were ties among some species that prevented their ordering along individual, unique past and future-predicted extinction sequences. To generate an observed extinction sequence, we broke ties as follows. For species with overlapping ranges, the species with the earliest year was ordered first. If the earliest years were the same, then we ordered the species with the earliest oldest bound first. For example, for a species with a range of 150–350 and one with a range of 150–200, we would order the second species before the first species. Finally, for species whose extinction year ranges were identical, we heuristically sampled ties in the extinction sequence and calculated the mean Δμ and Δσ^2^ across 10,000 tie-sampling replicates for empirical extinction. Notably, the species that lacked ties remain unchanged with this heuristic approach.

#### Extinction-Associated Traits

Our assessment of non-random extinction for *Leiocephalus* focused on morphological traits, because lizards have demonstrated a strong link between morphology and ecology (Losos 1990; Aerts et al. 2000; Melville and Swain 2000; Reilly et al. 2007; Goodman et al. 2008; Tulli et al. 2009; Edwards et al. 2022), and lizard ecology has been shown to explain patterns of extinction (Aerts et al. 2000; Reilly et al. 2007). For *Leiocephalus*, body size was the only trait for which data could be obtained for all 32 species. We also measured morphological diversity as body size-corrected morphological diversity for available extant species (*n* = 2 2 species). The difference in species coverage for morphological diversity data can be attributed to several species that lack suitable available vouchered museum specimens (e.g., species only known from fossils). Body size was measured as the maximum observed snout-vent length (SVL), a common measure of lizard body size that is independent of tail length. We obtained maximum SVL measures from an updated list of previous publications (Pregill 1981; Henderson and Powell 2009; Kemp and Hadly 2015; Köhler et al. 2016), except for *L. varius*, for which we obtained a maximum value from museum specimens.

To summarize extant morphological diversity, a principal component analysis (PCA) of 15 body size-corrected morphological characters was conducted. We measured fluid-preserved specimens for: snout-vent length, fore-hindlimb distance, pelvis width, pelvis height, total tail length, head length, head width, head height, finger IV metatarsal length, upper arm length, forearm length, toe IV metatarsal length, toe IV width, thigh length, and shank length. Prior to conducting the PCA, we normalized our data via log-transformation and then corrected for body size allometry by dividing trait values by snout-vent length (except snout-vent length). We then used the first two principal component axes, which jointly explained 69.3% of the variance, each as a trait that we tested for evidence of non-random extinction.

#### Extinction Model Selection

We tested four extinction sequences for evidence of non-random extinction under the three general models: past body size, predicted-future body size, and the predicted-future sequences of the two morphological diversity PCA axes. For body size extinction sequences, we evaluated all recorded extinctions starting with all described *Leiocephalus* species (*n* = 32, *e* = 8) and predicted-future extinctions starting with all extant species until only IUCN red list LC species remain (*n* = 2 4, *e* = 10). Similarly, morphological diversity predicted-future extinction sequences began with all extant species and ended with only LC species (*n* = 2 2, *e* = 8, based on available trait data). For each sequence, we separately evaluated Δμ and Δσ^2^ for each of the 10,000 tie-sampling replicates. There was little difference across replicates (Supplementary Figures 1,2), and so we calculated Δμ and Δσ^2^ at each time step across the tie-replicates for each of the four extinction sequences, and tested for non-random extinction in these mean extinction sequences.

### Testing Non-random Extinction Models

For a given data set of *n* species whose extinction sequence of *e* extinctions has been observed with respect to a trait *x*, simulations can be performed to ask if any of the three general models of non-random extinction fit the data better than a simple random extinction model. To do this, we developed simulations under each model assuming Gaussian distributions of continuous values for 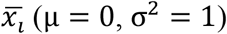, each with *n* species, the number in the observed data. Species were removed sequentially from the simulation, and Δμ and Δσ^2^ recorded at each loss step compared to starting μ and σ^2^ (up to the total number of extinctions observed in the data, *e*). For the random extinction simulation, the extinction sequence up to *e* proceeded independent of of *x* via random uniform sampling across the *n* species. We created two directional extinction simulations to test in which tail extinctions occur—one where the largest 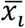 values are lost in decreasing order and another where the smallest are lost in increasing order. The simulation of disruptive extinction removed species with the smallest distances from μ in increasing order as calculated by the absolute value distance of each species mean from the clade-wide mean, 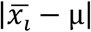. Lastly, simulation of stabilizing extinction used the same absolute value distance but removed species with the largest distances in decreasing order. For a given observed data set, each of the simulations were repeated 10,000 times. All R code functions used to perform these simulations can be found at https://github.com/ieco-lab/leiocephalus and can be adapted for any observed data set.

Once a distribution of simulations is produced under each of the extinction models, the observed data can be compared using a root-mean-square error (RMSE) goodness-of-fit statistic. For each extinction model, we first determined the mean values of Δμ and Δσ^2^ (μ_*sim*−*mean*_ and μ_*sim*−*variance*_) across the 10,000 simulations at each sequence step a species is lost relative to starting values. *μ*_*sim*−*mean*_ and *μ*_*sim*−*variance*_ were then used to calculate RMSE for each extinction model and species loss simulation (*RMSE*_*sim*−*mean*_ and *RMSE*_*sim*−*variance*_), as in (4) for Δμ and Δσ^2^ by replacing Δμ_*i*_ with 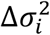 and μ_*sim*−*mean*_ with μ_*sim*−*variance*_.

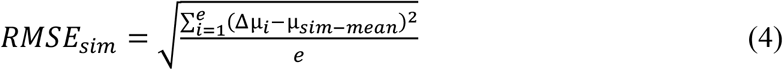

RMSE for observed extinction data relative to each extinction model was calculated with (4) by replacing simulated Δμ_*i*_ with the observed equivalent, producing five total sets of *RMSE*_*obs*−*mean*_ and *RMSE*_*obs*−*variance*_.

To conduct model selection for extinction models that jointly considers changes in both distribution metrics, we calculated Euclidean distances for each extinction model on a plane of Δμ and Δσ^2^. Before measuring distances, RMSE values were centered and rescaled. Mean values across simulations were then calculated for rescaled *RMSE*_*sim*−*mean*_ and *RMSE*_*sim*−*variance*_ for each of the five extinction models (μ_*RMSE*−*sim*−*mean*_ and μ_*RMSE*−*sim*−*variance*_). Euclidean distances were then measured between each of the five sets of μ_*RMSE*−*sim*_ points and (i) rescaled *RMSE*_*obs*_ to obtain a goodness-of-fit metric for the observed data and (ii) rescaled *RMSE*_*sim*_ to develop reference distributions for each extinction model. To assess significance for this goodness-of-fit metric and identify the best fit model, we conducted a nonparametric one-tailed Wilcoxon signed-rank test, W, comparing the Euclidean distances for each model. The Wilcoxon signed-rank test was used here, since Euclidean distances for all extinction models demonstrated a consistent right skew, which is suggestive of deviation from Gaussian distribution expectations that are necessary for analogous parametric tests like the z-test. Wilcoxon signed-rank tests were conducted to identify the extinction model with the highest W that also failed to reject the left tail null hypothesis, meaning that the simulated extinction model could have the same or greater Euclidean distance as the observed data and therefore could have come from the same distribution.

## Results

Past extinctions of *Leiocephalus* were best explained by directional extinction of species with large body size (Supplementary Figure 3, Figure 2). Model selection confirmed that directional extinction of large species was the only model that fit observed data significantly (Table 1, Supplementary Figures 4,5). In contrast, none of the extinction models fit the observed predicted-future extinction sequence for body size (Table 1, Supplementary Figures 4,5). The two best-fit models were the non-random model, where future loss was independent of body size, followed by the stabilizing model, where species of intermediate body size are the most likely to persist in the future. However, the observed data differed significantly from both models, indicating the possibility of a more complex extinction scenario that was not simulated under the three general models.

**Figure 2:**
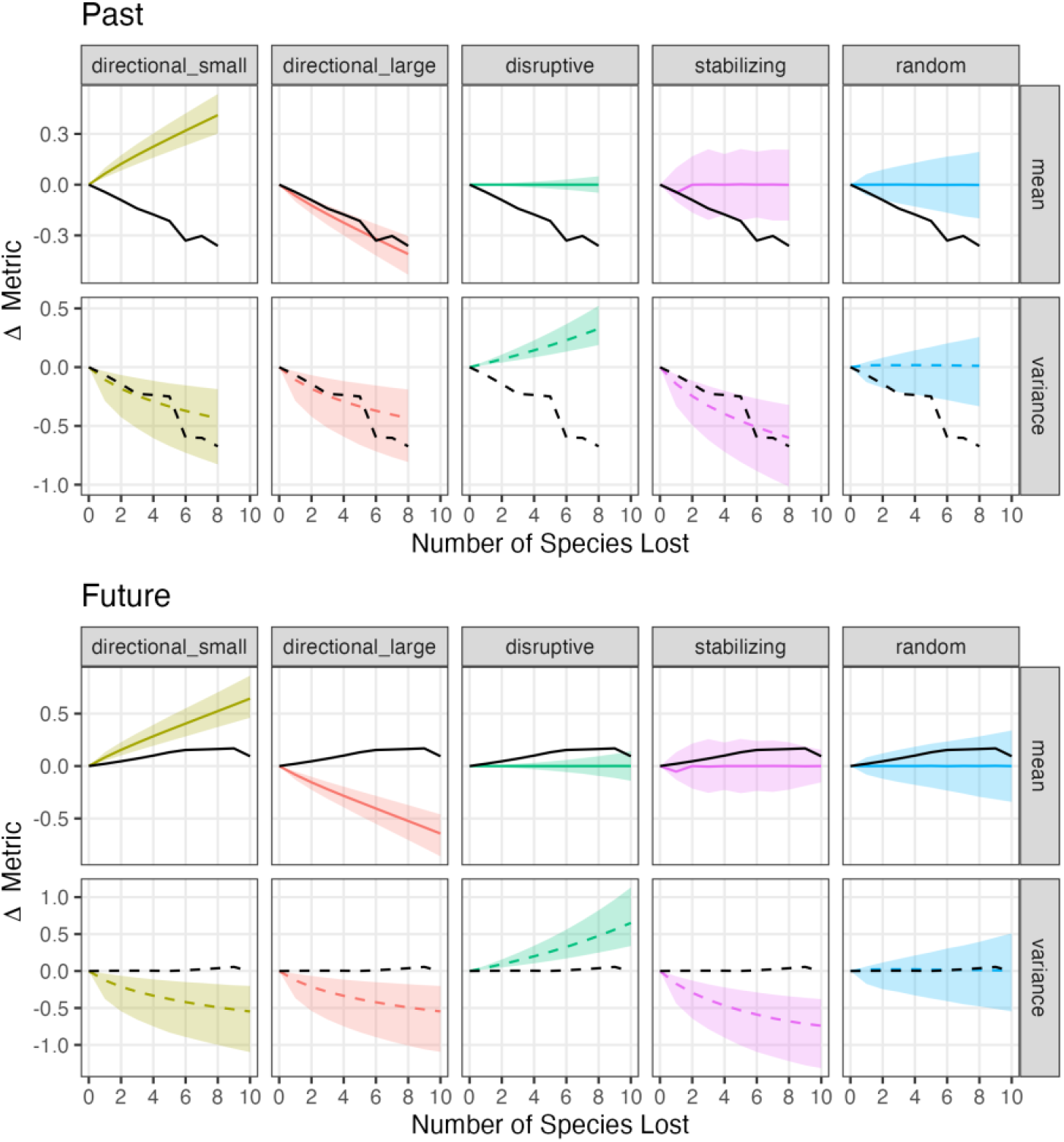
Extinction of body size through time for *Leiocephalus*. Species of *Leiocephalus* show directional extinction of large values in the past (TOP) and random extinction for predicted-future loss (BOTTOM) for body size as measured by maximum snout-vent length. For each analysis of extinction, changes in the clade-wide trait mean and variance are plotted as species are lost for the observed data (black solid and dashed lines) and for a distribution of simulations of the same species richness and number of extinctions for each extinction model. For each extinction model, median values are plotted as a solid or dashed line and the 95% confidence interval as the corresponding shaded region. The best fit extinction model is the one that observed data lines fall within the model distribution for changes in both mean and variance.

**Table 1:** Extinction model fit for *Leiocephalus* loss through time. Extinction models were fit for body size as measured by maximum observed snout-vent length and the first two axes from a morphological diversity PCA conducted on 15 traits. Significant model fits are presented in bold and are followed by an asterisk based on a one sample Wilcoxon signed-rank test.

| Trait | Timeframe | Model | W statistic |
| --- | --- | --- | --- |
| Body Size | Past | **Directional (large)** | **4.29E+07\*** |
|  |  | Stabilizing | 1.38E+05 |
|  |  | Random | 2.10E+01 |
|  |  | Directional (small) | 0.00E+00 |
|  |  | Disruptive | 0.00E+00 |
|  | Future | Random | 1.33E+07 |
|  |  | Stabilizing | 6.81E+04 |
|  |  | Directional (small) | 2.78E+02 |
|  |  | Disruptive | 4.20E+01 |
|  |  | Directional (large) | 0.00E+00 |
| Morphological PC1 | Future | **Random** | **3.75E+07\*** |
|  |  | Stabilizing | 3.35E+05 |
|  |  | Directional (small) | 4.40E+03 |
|  |  | Disruptive | 2.10E+01 |
|  |  | Directional (large) | 0.00E+00 |
| Morphological PC2 | Future | **Stabilizing** | **3.35E+07\*** |
|  |  | Random | 7.39E+05 |
|  |  | Directional (large) | 5.02E+03 |
|  |  | Directional (small) | 0.00E+00 |
|  |  | Disruptive | 0.00E+00 |

Further analysis of extinction for other morphological traits was first done by performing PCA on body-size corrected morphological traits. The first two principal component axes (PCs) accounted for 49.43% and 19.87% of explained variance (all subsequent PCs explained <10% of variance each), with trait loadings that suggested two axes that were describing different morphological features (Figure 4, Supplementary Table 1). For PC1, body size, pelvis width, head height, head length, head width, finger IV length, toe IV length, and toe IV width and for PC2, fore–hind limb distance, upperarm length, forearm length, thigh length, shank length, and total tail length all loaded strongly and positively for their respective PCs (Supplementary Table 1). Thus, we viewed PC1 as a composite measure of several traits that describe head, body, and digit size while viewing PC2 as a summary of limb and tail lengths. For the morphological PCA, the area of the minimum convex polygons occupied by all extant species was compared to that for least concern species only, showing that the polygon for least concern species occupies only 44.07% of the area for all extant species (Figure 4). Contraction from loss of threatened species is especially noticeable along PC2. Plots of predicted-future extinction for PC1 and PC2 yielded different trajectories (Figure 3). Empirical extinctions for PC1 were most similar to random extinction while those for PC2 were similar to stabilizing extinction. Extinction model selection confirmed that only random extinction significantly fit PC1 and only stabilizing extinction significantly fit PC2 (Table 1, Supplementary Figures 6,7).

**Figure 3:**
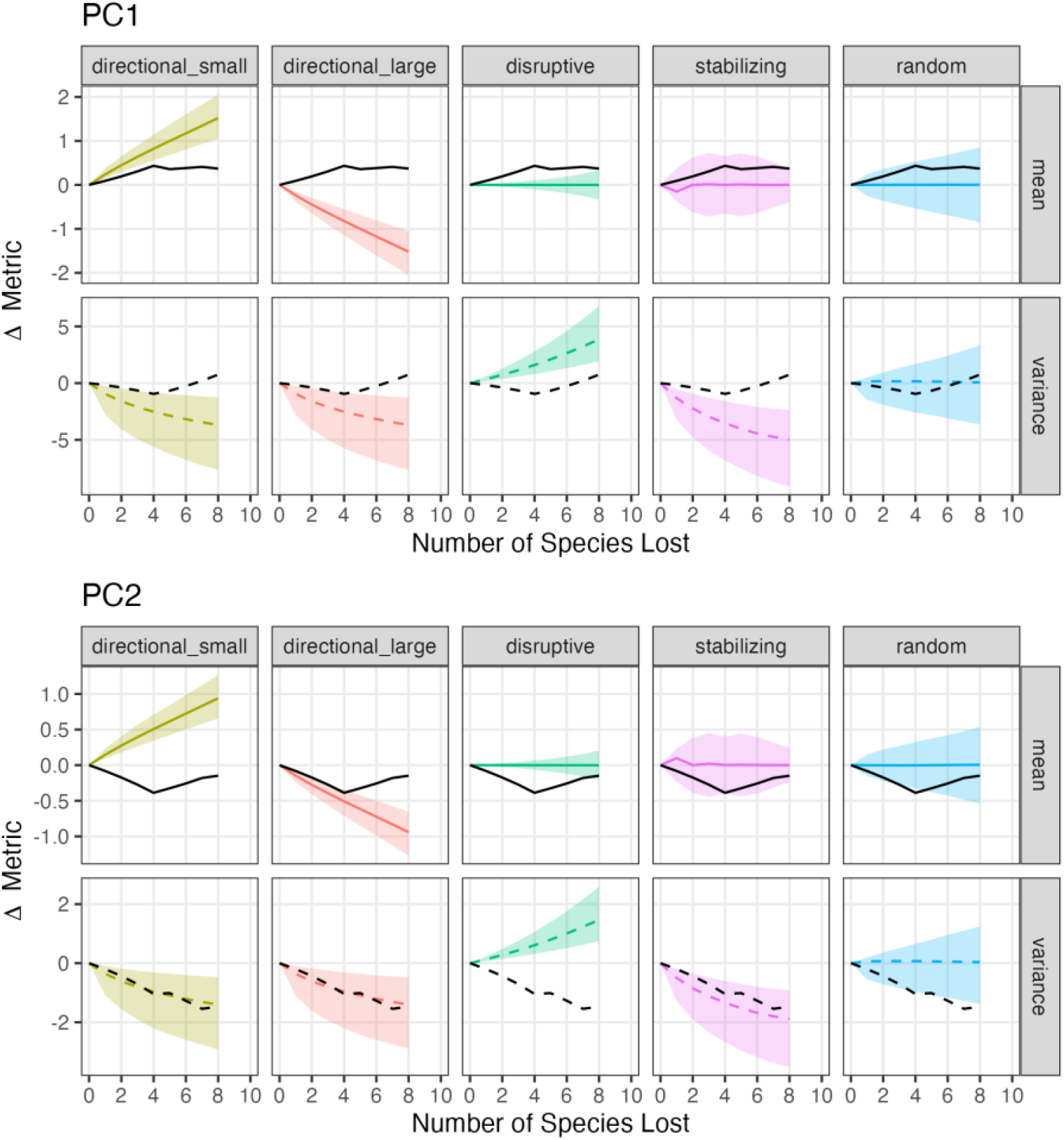
Future extinction of morphological diversity through time for *Leiocephalus*. For morphological diversity as measured by the first two axes of a PCA conducted on 15 traits, *Leiocephalus* predicted-future extinctions support two different patterns. The first axis (PC1) corresponds with traits that measure general body shape and size and is best explained by random extinction (TOP) while the second axis (PC2) describes appendage lengths and shows stabilizing extinction (BOTTOM). For each analysis of extinction, changes in the clade-wide trait mean and variance are plotted as species are lost for the observed data (black solid and dashed lines) and for a distribution of simulations of the same species richness and number of extinctions for each extinction model. For each extinction model, median values are plotted as a solid or dashed line and the 95% confidence interval as the corresponding shaded region. The best fit extinction model is the one that observed data lines fall within the model distribution for changes in both mean and variance.

**Figure 4:**
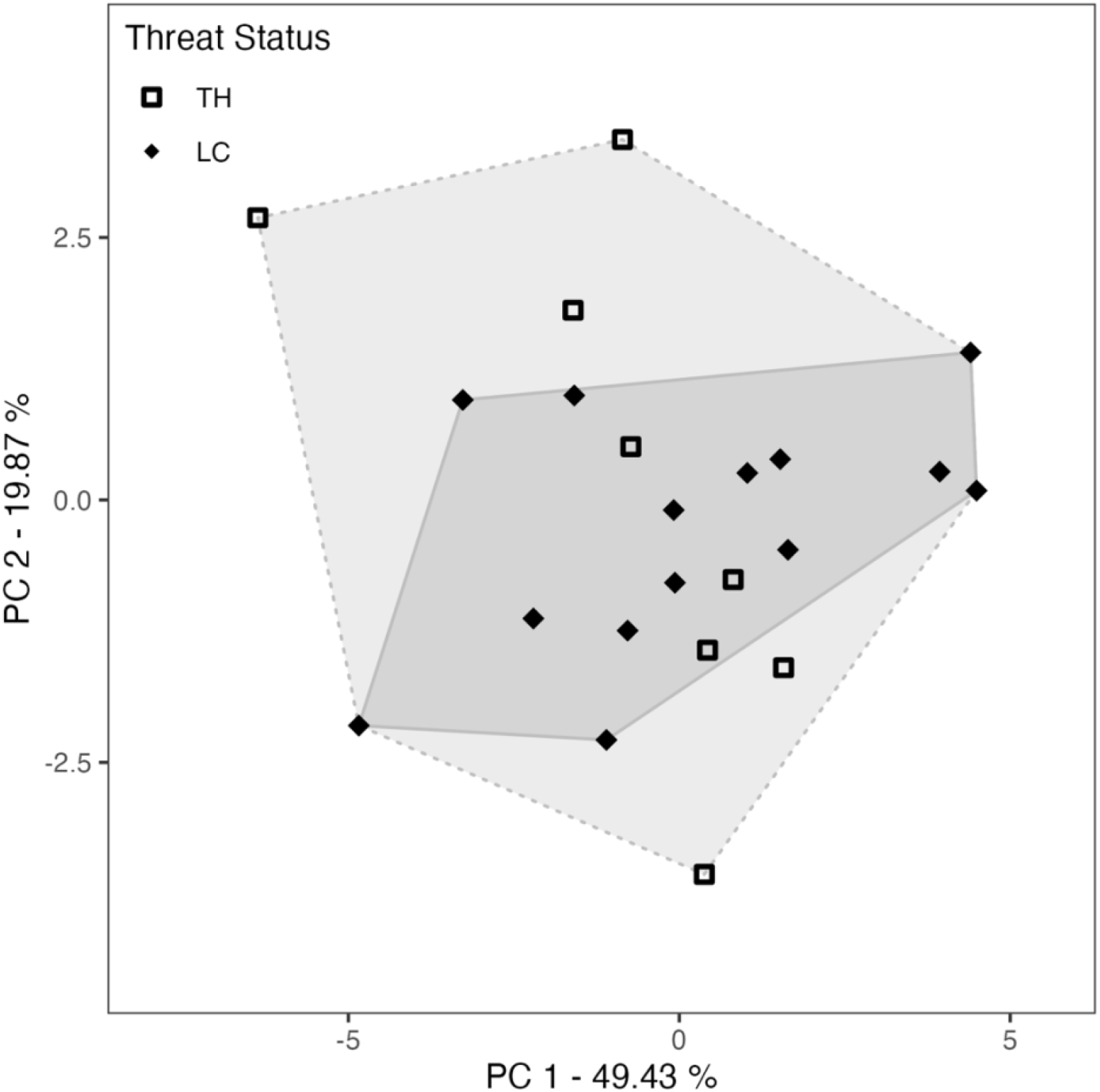
Morphological diversity principal components analysis (PCA). Predicted-future extinctions of *Leiocephalus* show a loss of morphological diversity as measured with PCA for 15 traits. The first two axes explained 59.30% of the variance and corresponded to general body shape and size (PC1) and appendage lengths (PC2). The minimum convex polygons for extant species (dashed line and lighter grey fill) and predicted-future species after extinctions expected from IUCN red list status (IUCN 2020) (solid line and darker grey fill) show considerable contraction, especially along PC2. Species are represented by black points that are shaped according to their grouped IUCN status.

## Discussion

As an imperiled clade, lizards in the genus *Leiocephalus* demonstrate changing non-random extinction for morphology. *Leiocephalus* extinctions to date appear driven by competition and predation by introduced mammals and hunting by humans, leading larger species to go extinct first (directional extinction; Borroto-Páez and Woods 2012; Kemp and Hadly 2015). For example, one of the largest species recorded, *Leiocephalus cuneus* was estimated at 200 mm SVL and was last seen in the 17th Century on Guadeloupe but previously was known from at least three other islands (Anguilla, Antigua, and Barbuda; Henderson and Powell 2009; IUCN 2020). The last observations of *L. cuneus* correspond with the arrival of European settlers who brought rodents, cats, dogs, and other mammals with them (Steadman et al. 1984), which have since been identified as contributing to extinction of native Caribbean faunas (Borroto-Páez and Woods 2012). Although direct evidence does not exist for the cause of all extinctions of *Leiocephalus*, examples like *L. cuneus* support the proposed drivers above and add to growing global trends (Sax and Gaines 2008; Bellard et al. 2016; Wood et al. 2017). Loss of large species has likely impacted Caribbean ecological communities. *Leiocephalus* are omnivorous, but diets differ across species (Schoener et al. 1982; Henderson and Powell 2009). Although many *Leiocephalus* tend to eat mostly invertebrates, some are saurophagous, feeding on other lizards like *Anolis* and have been shown to exert selective pressure on fitness-related traits in *Anolis* (Schoener et al. 2002; Henderson and Powell 2009; Vanhooydonck et al. 2009). Future extinctions stand to alter Caribbean ecosystems further but likely will differ in effect because of predicted non-random extinctions.

Predicted-future *Leiocephalus* extinctions are expected to be complex for body size with none of the general models fitting the observed data significantly, but with random and stabilizing models fitting the data best. Similarly, predicted-future extinctions demonstrate support for stabilizing extinction of *Leiocephalus* morphological diversity. This means that species with particularly small and large morphological trait values (when body size is accounted for) will be preferentially lost. In particular, species with extreme appendage lengths are expected to go extinct first. Loss of species with extreme values for these traits in *Leiocephalus* is noteworthy because as their common name suggests, they lift and curl their tails upwards for a number of reasons, including courtship and as a predator deterrent (Cooper 2001, 2007; Kircher and Johnson 2017). Tail curling in response to predation threat has been hypothesized to demonstrate fitness of the *Leiocephalus* individual and may be used for deterrence or deflection (and distraction via tail autotomy) of predatory attacks (Cooper 2007; Kircher and Johnson 2017). Stabilizing extinction here may reflect loss of species that are unable to optimize this anti-predator behavior because of short relative tail length or for whom failed deterrence may become too costly because of reduced fitness associated with autotomy of longer tails in many lizards (McElroy and Bergmann 2013). Similar stabilizing extinction of extreme limb lengths in *Leiocephalus* may reflect loss of species with more specialized locomotor performance on particular substrates (e.g., longer relative limb lengths performing better on firmer substrates like rocks and shorter on more fluid substrates like sand; Irschick and Garland 2001; Goodman et al. 2008) or association with particular habitats (e.g., Losos 1990; Melville and Swain 2000; Wiens et al. 2006; Losos 2009; Tulli et al. 2009; Edwards et al. 2022).

The transition from directional to stabilizing extinction for *Leiocephalus* may coincide with increased anthropogenic extinction pressures like habitat loss due to urban and agricultural development, charcoal production, and wood harvesting (Brooks et al. 2002; Zachos and Habel 2011; Leisher et al. 2013; IUCN 2020) as well as intensifying predation by invasive mammals like the mongoose, which can inhabit a wide range of habitats (Borroto-Páez and Woods 2012; IUCN 2020). Such pressures are great for Hispaniola, the most species rich island for extant *Leiocephalus* (Hedges 2022), where deforestation and development have become prolific over the last few decades (Sangermano et al. 2015; Hedges et al. 2018). Correspondence between habitat loss and extinction of species with extreme appendage lengths may result from loss of species that are more specialized to particular microhabitats (Blackburn and Gaston 2003). Additionally, invasive predators like the mongoose have now even penetrated remote mountains on islands like Hispaniola and prey readily on diurnal, ground-dwelling lizards like many species of *Leiocephalus* (Borroto-Páez and Woods 2012; Hedges and Conn 2012). Therefore, mongoose might indirectly benefit from the mismatch between more developed, less diverse habitats and *Leiocephalus* with more extreme phenotypes that are poorly suited to remaining habitats. Surviving *Leiocephalus* species then may be “microhabitat generalists” (Arnold 1998; Tulli et al. 2012) that perform at an intermediate level across different microhabitats and consequently manage to avoid predation.

Changing extinction pressures and the transition from directional to stabilizing extinction has conservation implications for *Leiocephalus*. Species prefer mesic and xeric habitats but habitats can range from coastal rocky shrubland and beaches to forests further inland (Henderson and Powell 2009). Regardless of type, some *Leiocephalus* have restricted habitat preferences and are therefore likely to be impacted as human development continues. Therefore, the diverse habitats of these species need to be protected. We recommend that development plans include setting aside resources for dedicated protected areas and reserves to maintain such habitat (Hedges et al. 2018). Similarly, we urge that control plans be formed for invasive species like the mongoose, which have already demonstrated capacity to cause Caribbean lizard extinctions (Hedges and Conn 2012). These efforts are especially important for conservation of *Leiocephalus*, because under our predicted stabilizing extinction, we would expect a reduction in *Leiocephalus* trait diversity, which will further diminish the clade’s ability to respond to future disturbances.

Shifts in non-random extinction for *Leiocephalus* can be understood as a conceptual extension of G.G. Simpson’s adaptive landscape (Simpson 1944). The adaptive landscape depicts a surface of fitness for all combinations of trait values and includes at least one region where a set of trait values maximize fitness, the adaptive peak (Arnold et al. 2001). For whole clades, species reside on the landscape and change in position to maximize viability by moving towards the nearest adaptive peak (Hansen 1997; Arnold et al. 2001). However, the surface can change, such as in response to altered environmental conditions or disturbances (Arnold et al. 2001). Species can respond to peak shifts, but those that fail to do so can go extinct. In the case of *Leiocephalus*, the past adaptive peak corresponded with a smaller body size, and species closer to that peak may have been able to avoid competition or predation from introduced mammals and hunting by humans because of it. Conversely, larger species were unable to respond quickly enough and went extinct. Over time, extinction pressures have changed, causing intermediate appendage lengths to be most viable (a pattern supported by an estimated extant *Leiocephalus* adaptive landscape; Supplementary Text 1, Supplementary Figures 8,9). Patterns in the adaptive landscape for *Leiocephalus* differ from those expected for larger or especially diverse clades, where multiple adaptive optima might exist across the landscape. In such cases, it may become difficult to resolve non-random extinction due to competing signals corresponding to different peaks. It may then become necessary to restrict analyses to a single peak of interest first to disentangle patterns of extinction, as is the case for current *Leiocephalus*. As such, in the absence of intervention or rapid shifts in phenotype, stabilizing extinction will shape *Leiocephalus* trait space.

Future development and enactment of methods to characterize non-random extinction should account for several factors. The case study here included a random extinction model that was recovered as the best but not significant fit for the data. Therefore, it may be difficult to recover a specific model for random extinction and alternative estimations should be considered. Additionally, we were able to identify stabilizing extinction, but it may be difficult to distinguish between it and random extinction in some cases. This may be caused by identical expectations for μ and smaller changes in σ^2^ for sets of fewer extinctions that correspond with adjacent, wide confidence intervals for both stabilizing and random extinction models. Difficulty in identifying stabilizing and random extinction models contrasts with distinct expected patterns and narrower confidence intervals for directional and disruptive extinctions. Nevertheless, methods like those here should be assessed via simulation to determine the limitations in statistical power in clades with varying levels, timescales and intervals, and confidence in extinction history. Our method also assumes traits with a Gaussian distribution as implemented here, but future extensions should explore other distributional parameters as well. Ultimately, addressing such questions will ensure confidence in extinction model fit that can be compared across taxa.

Characterizing patterns of extinction remains an important goal to improve our understanding of biodiversity change, and the methods we describe advance efforts to do so. By looking at extinction patterns across the tree of life, broad patterns in the effects of different sources of extinction pressure can emerge. Knowledge of such patterns can be applied to predict future changes in biodiversity and inform conservation efforts. In doing so, we will be better prepared to preserve biodiversity in the Anthropocene.

