## Supplementary Information for "Detecting stabilizing, directional, and disruptive patterns of anthropogenic species loss with general models of nonrandom extinction"

**This PDF file includes:**

Supplementary Text 1
Supplementary Table 1
Supplementary Figure 1
Supplementary Figure 2
Supplementary Figure 3
Supplementary Figure 4
Supplementary Figure 5
Supplementary Figure 6
Supplementary Figure 7
Supplementary Figure 8
Supplementary Figure 9
Supplementary References

**Supplementary Text 1**: Analysis of *Leiocephalus* Adaptive Landscape

A theoretical development of the extinction model selection methods I introduced is the description of shifting nonrandom extinction as an extension of G.G. Simpson's adaptive landscape (Simpson 1944). Simpson's adaptive landscape is an impactful metaphor for biological diversification that describes a multidimensional surface of fitness for all possible trait value combinations for a lineage. Elevation (the height of the surface) for particular trait combinations corresponds with its adaptive fitness. At least one region on the adaptive landscape maximizes fitness, the adaptive peak (Arnold et al. 2001). When a clade is evaluated on an adaptive landscape, individual species reside on the landscape and move towards the nearest adaptive peak to maximize viability (Hansen 1997; Arnold et al. 2001). However, adaptive landscapes are not necessarily static and are likely to change in response to altered environmental conditions, thereby prompting new sets of traits that correspond to adaptive optima (Arnold et al. 2001). Examples of altered environmental conditions have been suggested to correspond with known sources of extinction pressure (Arnold et al. 2001), such as introduction of invasive species or sudden destruction of habitat (such as anthropogenic development; Barnosky et al. 2011; Ceballos et al. 2015). Under pressures that correspond with topographical shifts in the adaptive surface, species can respond, depending on their phenotypic plasticity or genetic variation that corresponds with particular trait values. As such, species that successfully survive shifts likely do so via changes in the frequency of traits that better correspond with the "new" adaptive peak(s). The mechanisms by which species trait frequencies might change rapidly in response to peak shifts are varied and beyond the scope of this study. However, species that are unable to respond quickly enough go extinct, which may provide a clearer picture of how an adaptive landscape has changed over time.

Given the shifting extinction pressures described for *Leiocephalus* (see main text), it seems prudent to characterize the adaptive landscape for this genus. To do so, I assessed the number of selective regimes for *Leiocephalus* morphology with SURFACE (Ingram and Mahler 2013). SURFACE is a method that uses phylogenetic relationships and continuous trait data to assess if different lineages in a clade have corresponding selective regimes via model fitting with AIC. Different selective regimes are sequentially added as Ornstein-Uhlenbeck process (OU) models (termed "Hansen models" for SURFACE; Ingram and Mahler 2013). OU processes have previously been pointed to as reasonable depictions of evolution towards adaptive optima

(Hansen 1997), and thus Hansen models are suitable for describing selective regimes. Sequentially added Hansen models are jointly considered on the phylogenetic tree topology and evaluated for model fit (“forward phase”). Following identification of the best fit forward phase model, a second phase attempts to reduce the number of distinct regimes by converting distinct forward phase regimes into convergent regimes and evaluating the change in model fit. This “backward phase” assesses all pairwise combinations of collapsed regimes iteratively to identify the best fit model of selective regimes.

To estimate the adaptive landscape for *Leiocephalus*, I used a phylogeny of inferred relationships among *Leiocephalus* species from cytochrome-*b* [Cyt-*b*] mtDNA sequence data (Hedges, unpublished data) and the morphological trait data used for PCA (main text). For the phylogeny, a maximum likelihood phylogeny was constructed from a concatenated sequence alignment for all available species (n = 19) in MEGA7 (Kumar et al. 2016). Sequences were partitioned by codon position (all three codon positions and noncoding), and nucleotide substitution was modeled with the Generalized Time Reversible model with a discrete gamma distribution for rate variation. To properly polarize the phylogeny, we used three outgroup squamates, *Anolis carolinensis, Iguana iguana,* and *Sceloporus occidentalis*. The tree with the highest log likelihood was then selected for subsequent analyses. The resultant phylogeny branch lengths were multiplied by a constant (29.6, the mean clade age (Kumar et al. 2017)) and pruned to include only extant *Leiocephalus* (Supplementary Figure 8). For traits, the same 15 morphological traits were used to conduct phylogenetic PCA (hereafter pPCA; Revell 2009). Prior to pPCA, I log-transformed morphological trait data and corrected for body size allometry via phylogenetic size-correction of snout-vent length (SVL) against all other traits with GLS regression (Revell 2009). The resultant residuals and log-transformed SVL were rescaled and centered to conduct pPCA. I found that for the pPCA of *Leiocephalus*, 77.44% of the variance was explained in the first 4 PCs (Supplementary Figure 9). These four PCs were thus treated as traits for analysis with SURFACE.

I conducted SURFACE in R (R Core Team 2020) ver. 4.0.2 with the package surface (Ingram and Mahler 2013) ver. 0.5 under default settings with the pruned *Leiocephalus* phylogeny and first four PCs from the corresponding pPCA. After conducting both phases of the analysis, two selective regimes could be recovered, including the regime for the whole tree (Supplementary Figure 9). The second selective regime corresponded to a single branch containing *Leiocephalus raviceps*, an IUCN least concern species (IUCN 2020) with moderate loadings in our original morphological PCA. Both regimes were necessarily recovered as non-convergent because of the lack of other peaks fit in the data. Given the paucity of recovered adaptive regimes (and therefore lack of peak shifts), I assessed the model fit of the SURFACE analysis against Brownian motion and a single peak OU process. Of the three models, Brownian motion fit the data least well (AICc = 326.918), whereas the single peak OU and SURFACE models better fit the data but had similar fits (AICc = 292.132 and 292.012, respectively). Given the similarity of the two best fit models, I plotted the peaks recovered for each (Supplementary Figure 9), which showed that the lineage-wide regime and OU peak to be in close proximity across PCs. Given the negligible differences in model fit for these two models and the principle of parsimony, it seems prudent to assume that based on the species and traits analyzed here, *Leiocephalus* likely have a single adaptive peak.

The analysis here suggests a single adaptive peak for extant *Leiocephalus* morphology, which is an unsurprising result. Recovery of an adaptive landscape peak shift corresponding to the shifting extinction pressures observed in the main text was unlikely, given the data that are currently available for *Leiocephalus*. Notably, analyses in the main text compared both extinct and extant species to suggest shifts linked to extinction. Phylogenetic relationships among several extant and all extinct species were not able to be reconstructed, as the inferred phylogeny was based solely on molecular data. Such molecular data was unavailable for missing species, either due to lack of compatible sequence data or material to sequence (several extinct species are known only from fossils, see main text). The case was the same for morphological trait data to a lesser degree. As such, part of the data necessary to detect the past extinction to expected future extinction shift were unavailable for this study. I suggest that future research should pursue the methods outlined here and in the main text for *Leiocephalus* and other imperiled clades, should sufficient data become available. Nevertheless, it appears that a single adaptive peak for extant *Leiocephalus* corresponds with an intermediate phenotype for the morphological traits considered here (Supplementary Figure 9), even though trait loadings for the morphological PCA and pPCA are similar but not identical. As such, the results of this analysis may be viewed as additional support for selection that corresponds with stabilizing extinction of morphological traits as described in the main text.

**Supplementary Table 1**: Axis loadings for morphological principal components analysis for *Leiocephalus*.

| Trait | Description | PC1 | PC2 |
|---|---|---|---|
| 1 | SVL | 0.3197 | 0.0031 |
| 3 | Fore-Hind Limb Distance | -0.1315 | 0.3489 |
| 6 | Pelvis Width | 0.3078 | -0.0974 |
| 7 | Pelvis Height | 0.2738 | -0.2651 |
| 9.1 | Total Tail Length | -0.1411 | 0.2399 |
| 11 | Head Length | 0.2638 | -0.0273 |
| 12 | Head Width | 0.3248 | -0.1566 |
| 13 | Head Height | 0.3091 | -0.1589 |
| 17 | Finger IV Metatarsal Length | 0.2975 | 0.1335 |
| 20 | Upperarm Length | 0.1518 | 0.4619 |
| 21 | Forearm Length | 0.1903 | 0.4265 |
| 23 | Toe IV Metatarsal Length | 0.2868 | 0.0026 |
| 24 | Toe IV Width | 0.2728 | -0.2138 |
| 26 | Thigh Length | 0.2660 | 0.2585 |
| 27 | Shank Length | 0.2124 | 0.4059 |

**Supplementary Figure 1**: Heuristic path of body size extinction. Heuristic resampling of extinction order for *Leiocephalus* body size extinctions shows little variation in changes to the clade-wide mean and variance for extinctions to date (lines in dark grey zone) and predicted-future extinctions of all but least concern species (lines in light grey zone). Sampling of least concern species (lines in white zone) shows an expected set of varying trajectories.

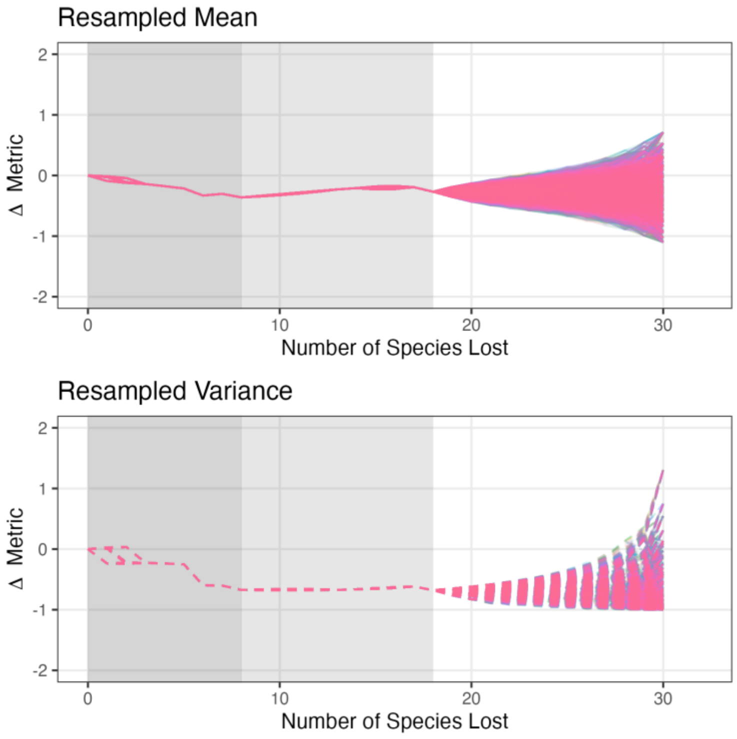

**Supplementary Figure 2**: Heuristic path of future morphological diversity extinction. Heuristic resampling of extinction order for *Leiocephalus* morphological diversity as measured by the first two axes in a principal components analysis (PCA) of 15 traits shows little variation in changes to the clade-wide mean and variance for predicted-future extinctions of all but least concern species (lines in light grey zone) for the first PC (LEFT) and second PC axis (RIGHT). Sampling of least concern species (lines in white zone) shows an expected set of varying trajectories.

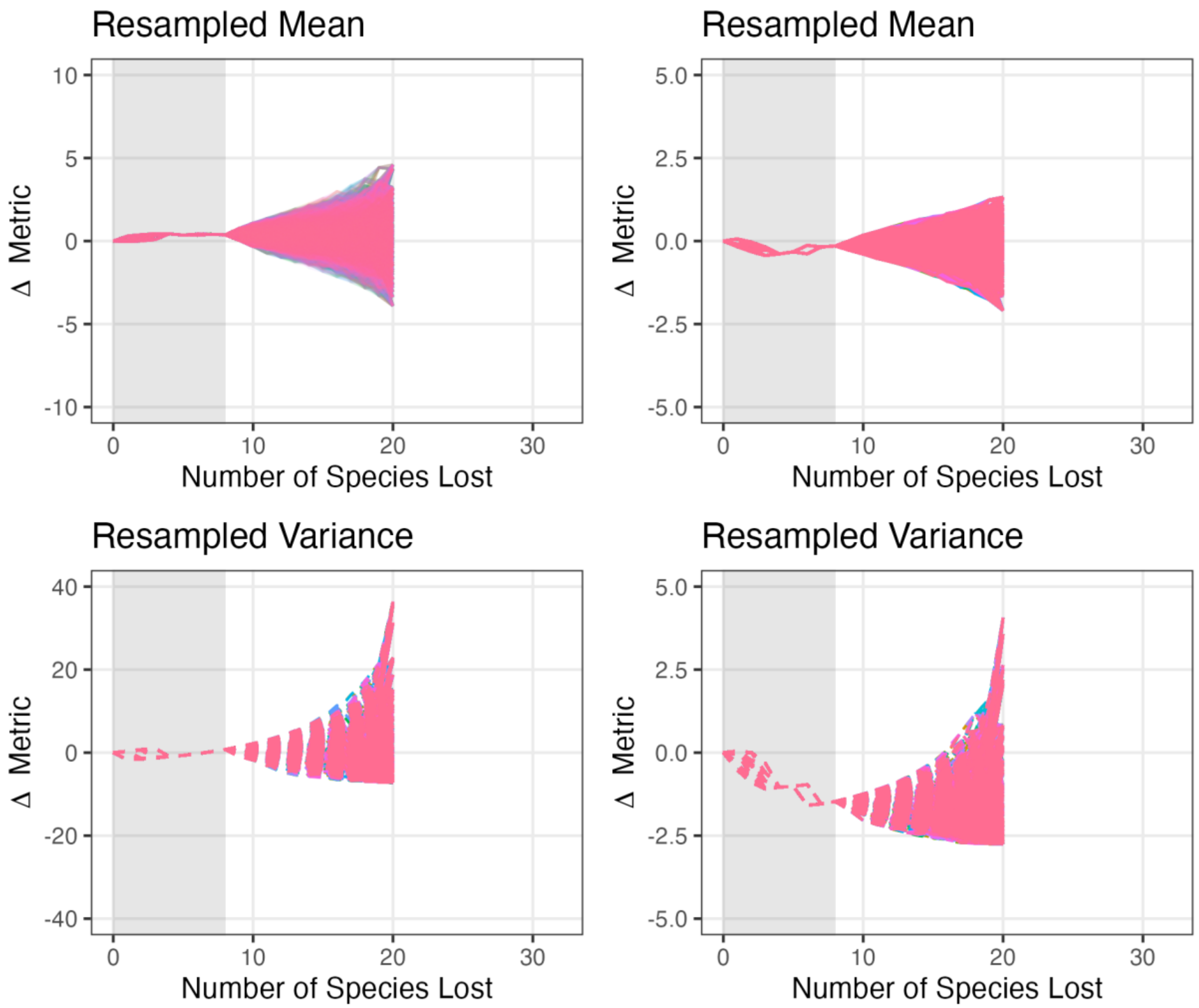

**Supplementary Figure 3**: *Leiocephalus* body size by grouped threat status. Extinction of *Leiocephalus* species to date seem to predominantly include larger species (EX). Extant species do not appear to differ in body size when compared by threat status (least concern species, LC, and all other threat categories as threatened, TH, based on IUCN red list status; IUCN 2020).

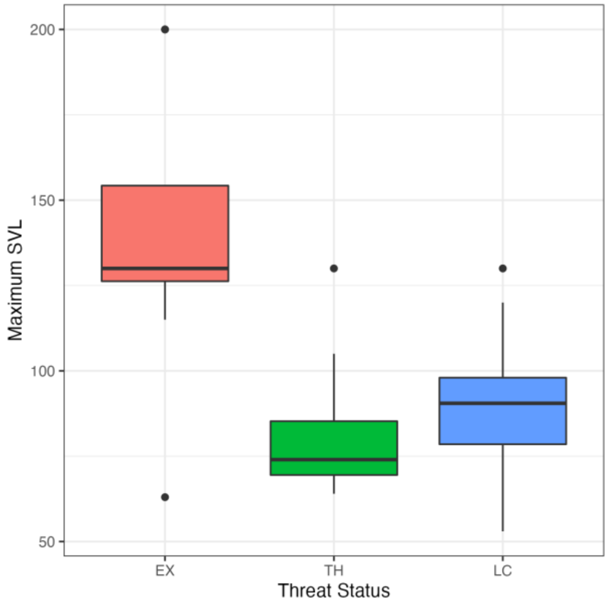

**Supplementary Figure 4**: Extinction of body size Euclidean space plot. Scatter plots of the scaled root mean square error (RMSE) for mean and variance on which Euclidean distances are calculated compare extinction models for body size. Fit for an extinction model is best when the observed data (black star) is closest the centroid for that model (black cross) and its corresponding simulations (colored points), which here are closest to directional extinction of large species for the past and random for predicted-future extinction with respect to body size for *Leiocephalus*.

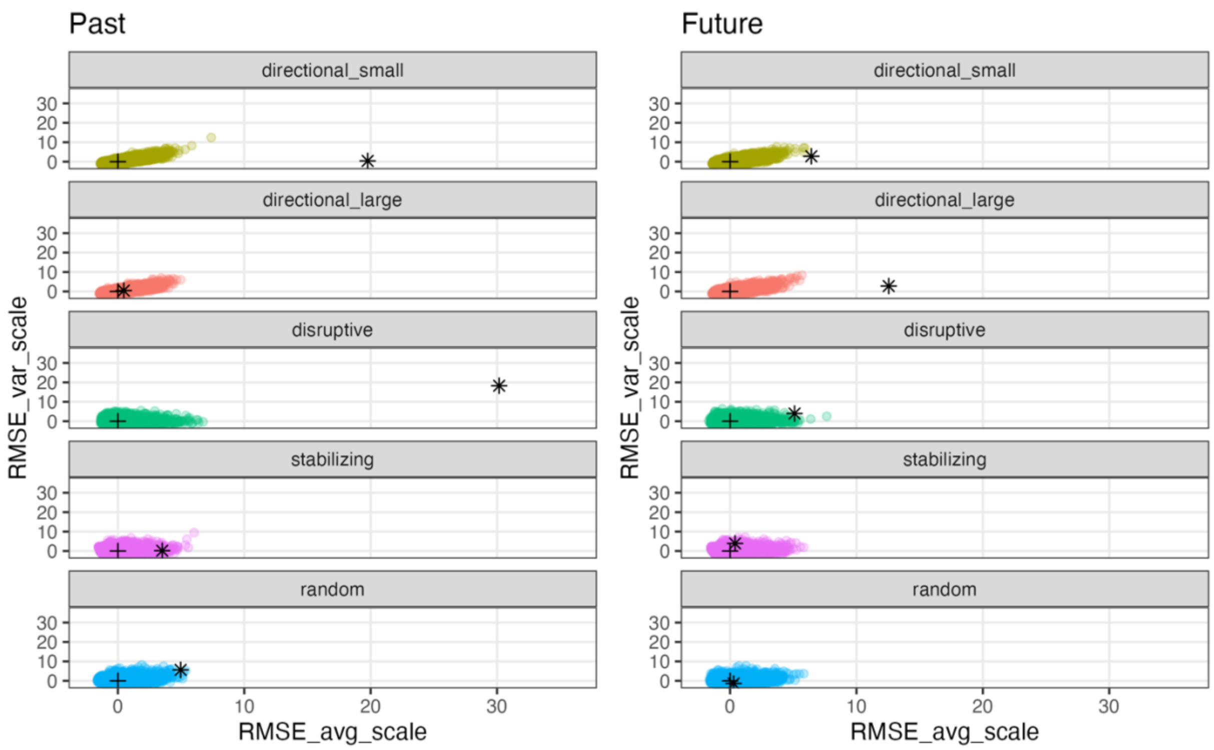

**Supplementary Figure 5**: Extinction of body size histogram. Histograms that compare extinction models for body size for goodness-of-fit based on Euclidean distances between observed data (vertical black line) and simulations (colored histograms) support past directional extinction of large species and future random extinction with respect to body size for *Leiocephalus*.

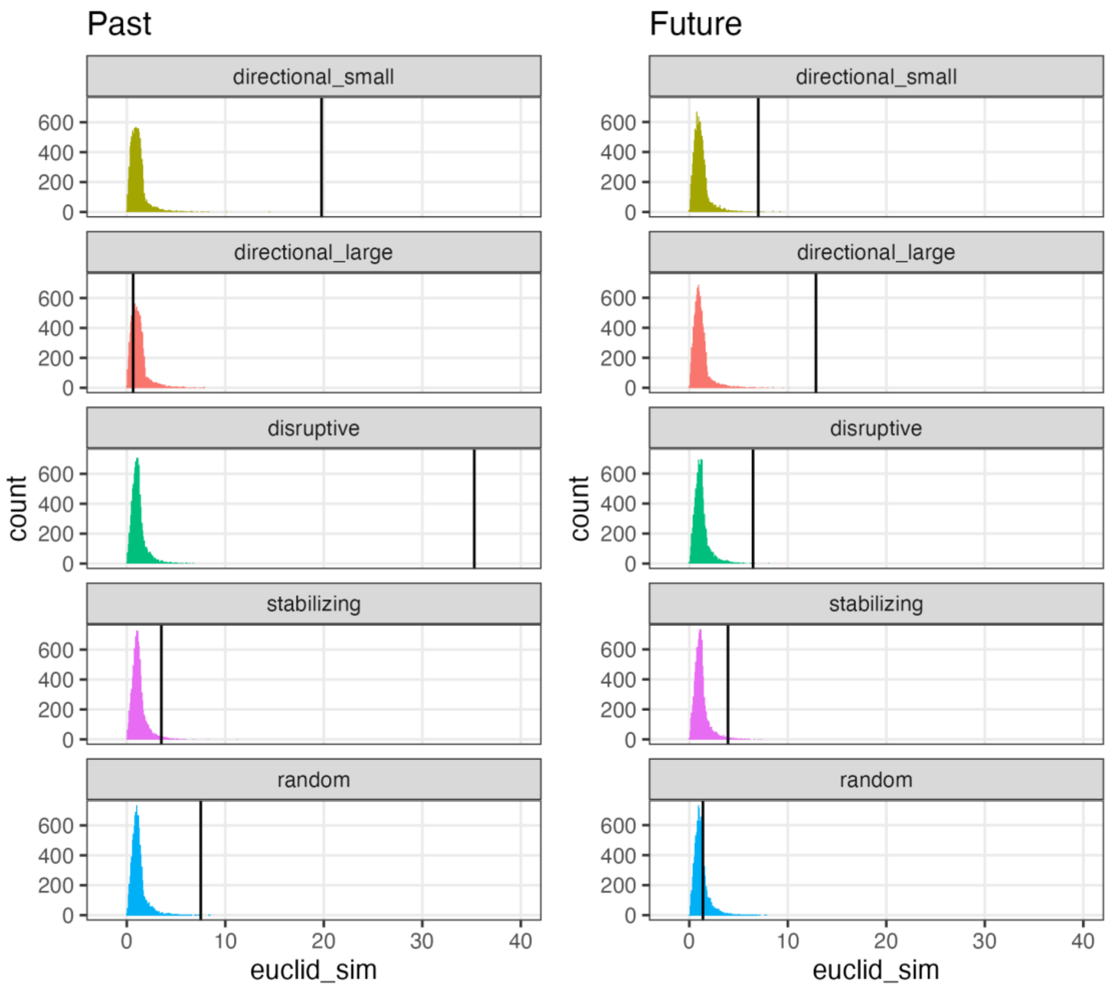

**Supplementary Figure 6**: Extinction of morphological diversity Euclidean space plot. Scatter plots of the scaled root mean square error (RMSE) for mean and variance on which Euclidean distances are calculated compare extinction models for morphological diversity as measured by the first two axes in a principal components analysis (PCA) conducted on 15 traits. Fit for an extinction model is best when the observed data (black star) is closest the centroid for that model (black cross) and its corresponding simulations (colored points), which here show predicted-future extinctions that are closest to random for the first axis (PC1) and stabilizing for the second (PC2) for *Leiocephalus*.

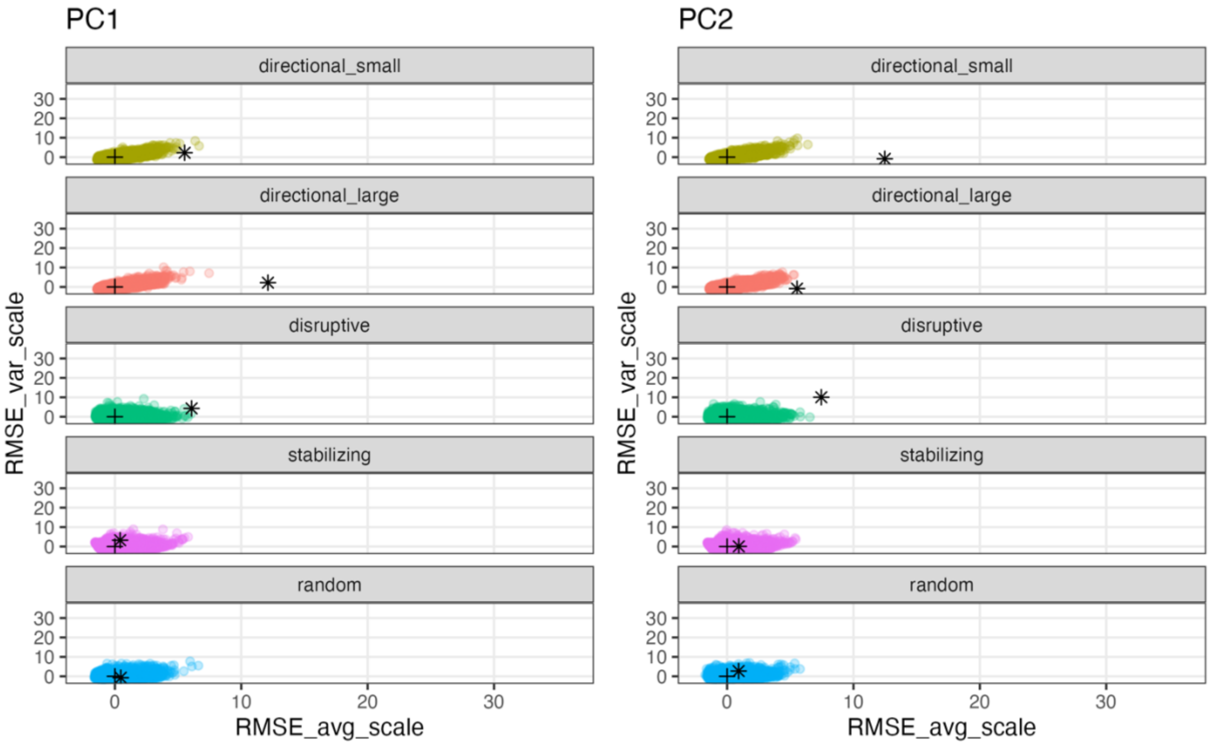

**Supplementary Figure 7**: Extinction of morphological diversity histogram. Histograms that compare extinction models for morphological diversity as measured by the first two axes in a principal components analysis (PCA) conducted on 15 traits for goodness-of-fit based on Euclidean distances between observed data (vertical black line) and simulations (colored histograms). Predicted-future extinctions are closest to random extinction for the first axis (PC1) and stabilizing extinction for the second (PC2) for *Leiocephalus*.

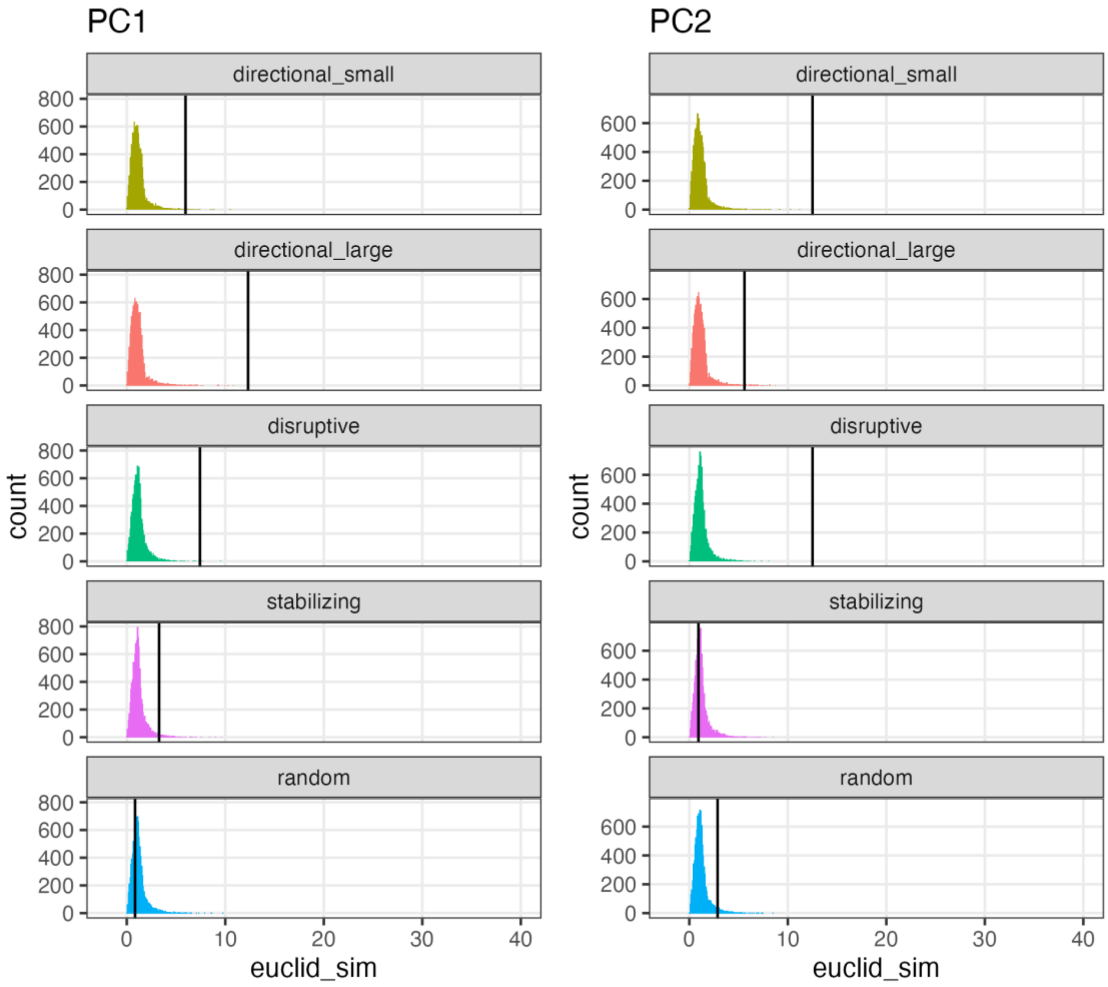

**Supplementary Figure 8**: Phylogeny of extant *Leiocephalus* used for SURFACE method. A maximum likelihood phylogeny constructed from molecular data shows the relationships among extant species of *Leiocephalus*. Tip labels depicted with red text correspond to species that are threatened and black text for those that are least concern according to the IUCN red list (IUCN 2020: all non-least concern species are TH and least concern species LC).

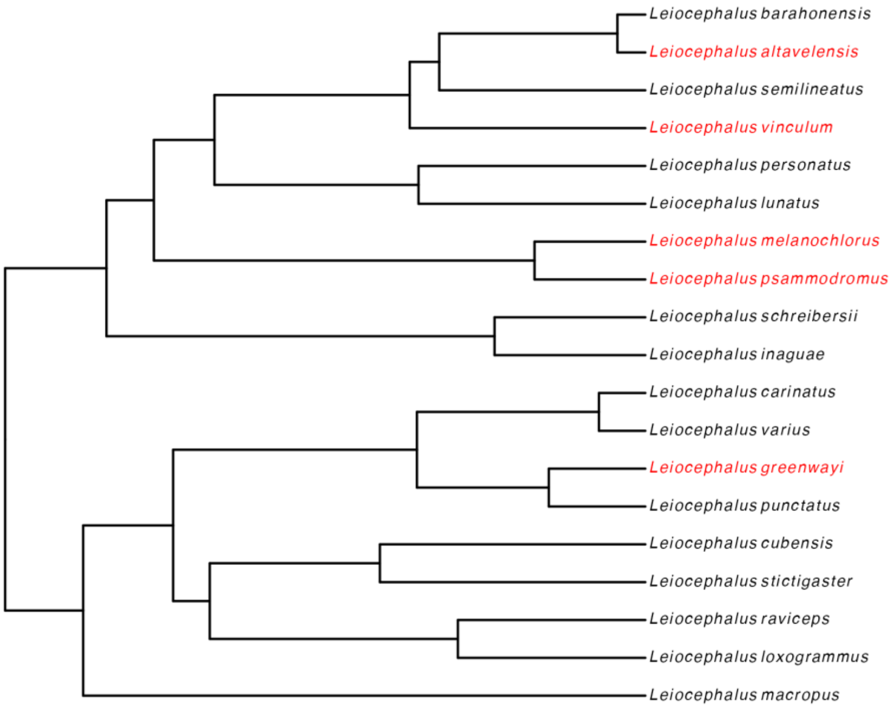

**Supplementary Figure 9**: Phylogenetic principal components analysis (PCA) of *Leiocephalus* with adaptive peaks. Phylogenetic PCA results are plotted for the first four PCs used in the SURFACE analysis. Individual species of *Leiocephalus* are represented by black points with point shapes that correspond to grouped IUCN red list status (IUCN 2020: all non-least concern species are TH and least concern species LC). Adaptive peaks recovered by SURFACE and single peak Ornstein-Uhlenbeck process models are represented by red points that are labeled. SURFACE label suffixes correspond to the whole-lineage peak (a) and single species peak for *Leiocephalus raviceps* (b).

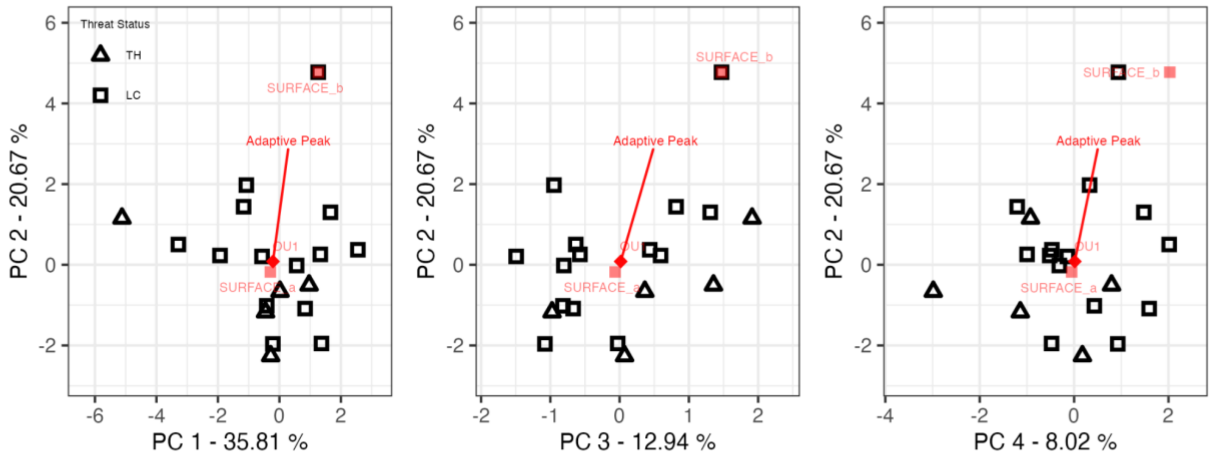
